# Long-insert sequence capture detects high copy numbers in a defence-related beta-glucosidase gene β*glu-1* with large variations in white spruce but not Norway spruce

**DOI:** 10.1101/2023.08.18.551884

**Authors:** Tin Hang Hung, Ernest T. Y. Wu, Pauls Zeltiņš, Āris Jansons, Aziz Ullah, Nadir Erbilgin, Joerg Bohlmann, Jean Bousquet, Inanc Birol, Sonya M. Clegg, John J. MacKay

**Affiliations:** Department of Biology, University of Oxford, Oxford OX1 3RB, United Kingdom; Latvian State Forest Research Institute “Silava”, LV-2169 Salaspils, Latvia; Department of Renewable Resources, University of Alberta, Edmonton, AB, T6G 2E3, Canada; Michael Smith Laboratories, University of British Columbia, Vancouver, BC, V6T 1Z4, Canada; Department of Botany, University of British Columbia, Vancouver, BC, V6T 1Z4, Canada; Department of Forest and Conservation Sciences, University of British Columbia, Vancouver, BC, V6T 1Z4, Canada; Canada Research Chair in Forest Genomics, Forest Research Centre, Université Laval, Québec, QC, G1V 0A6, Canada; Canada’s Michael Smith Genome Sciences Centre, BC Cancer Agency, Vancouver, BC, V5Z 4S6, Canada

**Keywords:** acetophenone pathway, targeted capture, *Picea*, CNV, secondary metabolism, conifer genomics

## Abstract

Conifers are long-lived and slow-evolving, thus requiring effective defences against their fast-evolving insect natural enemies. The copy number variation (CNV) of two key acetophenone biosynthesis genes *Ugt5*/*Ugt5b* and β*glu-1* may provide a plausible mechanism underlying the constitutively variable defence in white spruce (*Picea glauca*) against its primary defoliator, spruce budworm. This study develops a long-insert sequence capture probe set (Picea_hung_p1.0) for quantifying copy number of β*glu-1*-like, *Ugt5*-like genes and single-copy genes on 38 Norway spruce (*Picea abies*) and 40 *P. glauca* individuals from eight and nine provenances across Europe and North America respectively. We developed local assemblies (Piabi_c1.0 and Pigla_c.1.0), full-length transcriptomes (PIAB_v1 and PIGL_v1), and gene models to characterise the diversity of β*glu-1* and *Ugt5* genes. We observed very large copy numbers of β*glu-1*, with up to 381 copies in a single *P. glauca* individual. We observed among-provenance CNV of β*glu-1* in *P. glauca* but not *P. abies*. *Ugt5b* was predominantly single-copy in both species. This study generates critical hypotheses for testing the emergence and mechanism of extreme CNV, the dosage effect on phenotype, and the varying copy number of genes with the same pathway. We demonstrate new approaches to overcome experimental challenges in genomic research in conifer defences.

## Introduction

Conifers are keystone species in many terrestrial biomes^1^. They play an important role in global biogeochemical cycles, including the carbon, nutrient, and water cycles. They are also of large economic importance globally as a resource, derived from naturally regenerated forests and plantations, for timber and non-timber products^2^. However, forests and tree populations are facing threats from environmental change, most prominently from warmer climates^3^ as well as pest and pathogen outbreaks^4,5^. As part of their sessile lifestyle, longevity, and slow evolutionary response, conifers have developed effective defence systems against diverse faster-evolving natural enemies, which are key attributes to the forest health^6–8^.

One such defence system relies on acetophenones and their glucosides in white spruce (*Picea glauca*) across northern North America. Acetophenones and their glucosides are the most abundant soluble phenolic and glucosidic compounds in their shoots^9^. Variation in the constitutive accumulation of two particular acetophenones, piceol and pungenol, has been demonstrated to confer defensive properties and resistance against spruce budworm (*Choristoneura fumiferana*, Lepidoptera: Tortricidae), which is largely sympatric with white spruce^10^. Higher levels of these acetophenones reduce the budworm defoliation, which has been linked to decreased larval survival and pupal mass and delayed development in female moths, and increased fitness in the tree host^11^. Resistant white spruces have a high level of foliar piceol and pungenol, while both resistant and non-resistant trees can accumulate high levels of their glycosylated forms, picein and pungenin^12^. Phenology studies of white spruce and spruce budworm showed the peak of piceol and pungenol accumulation in hosts temporarily matches with the larval stage that is most damaging^10^. These acetophenones are conserved to some extent in Pinaceae and most *Picea* species accumulate at least one acetophenone glucoside in their foliage. Besides *P. glauca*, Norway spruce (*P. abies*), which is distributed across Europe, is the only spruce species known to accumulate both picein and pungenin^13^.

The biosynthesis and function of these acetophenones has been characterised by metabolite analyses, gene discovery and gene expression analysis, biochemical characterisation of biosynthetic enzymes, and insect *in vivo* assays. The formation of pungenin and isopungenin from pungenol is catalysed by UDP-sugar dependent glucosyltransferases, encoded by the *PgUgt5b* and *PgUgt5* genes respectively^14^, while genes and enzymes for formation of picein from piceol are still unknown. The glucosidic bond can be hydrolysed by a β-glucosidase encoded by the *Pg*β*glu-1* gene to release the active piceol and pungenol^15^. Substantial geographical and seasonal variations exist in the expression of *Pg*β*glu-1* linked to the phenotypic variation of acetophenone accumulation and tree survival^16^, with up to 1,000-fold difference in the gene expression between resistant and non-resistant trees.

However, the genetic underpinning of this phenotypic variation remains largely unknown and copy number variation (CNV) could be a plausible mechanism. Copy number variation was first studied in humans^17,18^ and was found to occur in 4.8–9% of the human genome, which is more frequent than single nucleotide polymorphisms^19,20^. There has been a lack of consensus on the definition of CNV, for example, the size of CNVs has been reported to range from 50 bp to several Mbps, but shorter ones are sometimes classified as indels^21^. CNV may be best defined as ‘the relative difference in copy numbers of particular DNA sequences among conspecific individuals^22^. CNV can lead to gene expression differences reaching several orders of magnitude^23^ brought about by various mechanisms, including simple dosage effects, changes in gene regulatory regions, and alterations to the physical proximity of genes in relation to their regulatory elements^24^.

Although CNV may explain phenotypic variation in important adaptive traits, there are very few studies that have investigated intraspecific or population level CNV in conifers. These studies have mostly focused on individual loci, such as the thaumatin-like protein gene in *Pinus sylvestris* (PsTLP)^25,26^ and the (+)-3-carene synthase gene in *Picea sitchensis* (PsTPS-3car)^27,28^ or mapped genomic CNV at the species level^29–31^. In *P. glauca* two genomic scans detected more than 1,300 and 3,900 CNVs, representing around 10% and 32% of the genes tested, respectively and, significant over-representation of defence response genes^32,33^.

The very large size of conifer genomes, between 18–35 Gbp for most species^34^, has raised evolutionary questions regarding their ability to adapt to changing biotic pressures and has also posed experimental limitations and challenges. While gene duplication has a central role in genome evolution, conifers generally retain more nonfunctional gene copies than other species and accumulate as high as 4% of gene-like sequences^35^. They also have a larger amount of intronic sequences containing more repetitive elements compared to flowering plants^36^. The abundance of pseudogenes also has challenged a contiguous assembly for their genomes and curation of a well-characterised gene catalogue^1^.

The sheer size and complexity of conifer genomes make whole genome sequencing impractical for studying conifer populations, therefore, CNV studies in conifer trees have utilised either array Comparative Genome Hybridization (aCGH) or quantitative PCR (qPCR). However, these methods have a significant limitation due to the lack of sequence data on the signals detected, which may result from non-targets, and unquantified variation among individuals in on-target signals. Alternatively, shotgun sequencing with short reads could confound non-functional gene-like sequences or pseudogenes from their highly similar parent genes^37^, and has substantial amplification bias such as GC content^38^. Therefore, a new method to bridge this gap in precision requires incorporating both long-read sequencing, which enhances the confidence of sequence validation, and targeted capture, which reduces the cost significantly for gigagenomes like spruces’, and makes experiments scalable for large population studies.

Here, we aim to develop a long-insert sequence capture protocol for quantifying copy number based on known gene targets in acetophenone biosynthesis in *P. glauca*. Our objectives are three-fold: (1) to design a probe set suitable across *Picea* spp. that captures the target genes β*glu-1* and *Ugt5*, which regulate the accumulation and release of acetophenone defence compounds, and single-copy genes as internal standards; (2) to characterise the identity and diversity of target gene models by integrating genomic sequence capture, transcriptome assemblies, and local gene assemblies; (3) to quantify the copy numbers of β*glu-1* and *Ugt5* and compare them among populations of *P. abies* and *P. glauca* using a coverage-based algorithm. The development of this protocol with the probe set particularly addresses the gaps in methods and resources in studying the genetic underpinning of insect resistance in conifer species.

## Methods

### Plant materials and nucleic acids extraction

Foliage samples for *Picea abies* and *Picea glauca* were collected in September 2021 from two common gardens located in Calling Lake, Alberta, Canada (55.28°, –113.17°), established in 1982^39^, and Saldus, Courland, Latvia (56.85°, 22.52°), established in 1975^40^ respectively. We collected foliage samples from the top-half crown of 38 and 40 individual trees, which represented 8 and 9 provenances respectively (**Figure 1** and **Supplementary Table 1**), compliant with relevant institutional guidelines and national legislations. Foliage samples were snap-frozen immediately after removal from the tree and stored at –80°C to preserve the integrity of nucleic acids until analyses.

**Figure 1.**
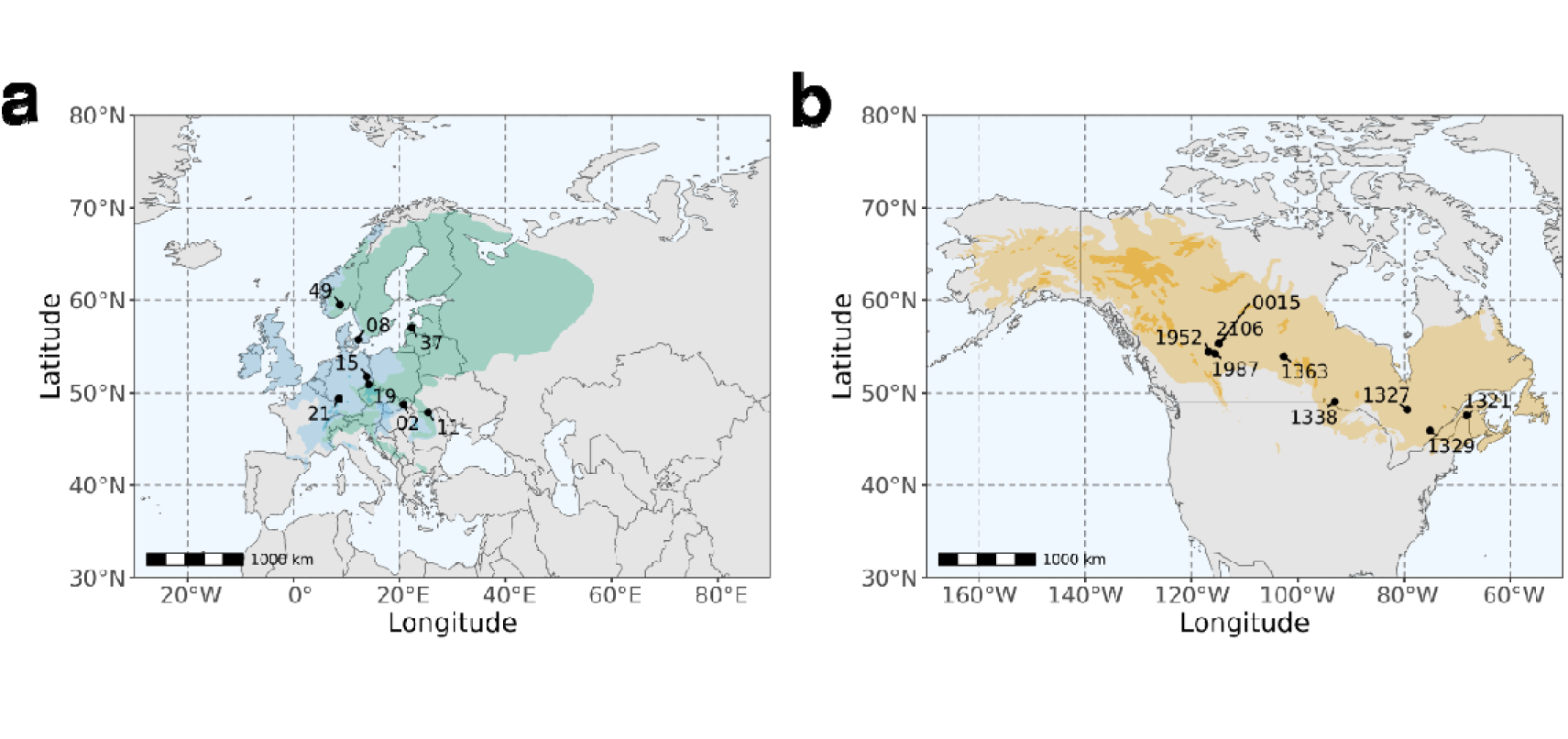
The localities of provenances (circles) of **(a)** P. abies and **(b)** P. glauca. For P. abies, the green polygon shows its native distribution, and the blue polygon shows its introduced and naturalised distribution. For P. glauca, the orange polygon shows its native distribution. Species distribution maps are obtained from ^87^ and ^88^ respectively.

We extracted genomic DNA and total RNA using the DNeasy Plant Mini Kit (Qiagen, United Kingdom) and the NEB Monarch Total RNA Miniprep Kit (#T2010, New England Biolabs, United Kingdom) respectively, determined their quantity on a Qubit 4 Fluorometer (Thermo Fisher Scientific, United Kingdom), and assessed their purity using a NanoDrop One Spectrophotometer (Thermo Fisher Scientific), with A260/280 and A260/230 above 1.80. Integrity was verified on a 0.5% agarose gel for genomic DNA and a 2100 Bioanalyzer (Agilent Technologies, United States) for total RNA.

### Selection of target regions and bait development

Full-length cDNA sequences of *Pg*β*glu-1* (NCBI GenBank Accession: KJ780719.1)^12^, *PgUgt5* (KY963363.1), and *PgUgt5b* (KY963364.1)^14^ were searched against the gene models of *P. glauca* WS77111_v2^2^ using BLAST 2.11.0+^41^. Their corresponding genomic coordinates were padded for 1,000 bp in both upstream and downstream directions, resulting in 24 target genomic sequences with a total length of 108,485 nt. We designed 80-nt baits at 2X tiling density to cover these sequences and produced a total of 2,427 baits. We filtered the baits according to number of BLAST hits against reference genome per bait and the predicted melting temperature between the baits and the BLAST hits, and retained 1,836 baits. We performed BUSCO analysis^42^ on the gene models to predict single-copy genes in the assemblies indicated above, 134 of which were randomly selected to serve as internal standards. Using the same design approach, we produced 21,951 baits. After filtering, 18,164 baits were retained to cover 913,726 nt. The final bait set covering acetophenone biosynthesis target genes and single-copy internal standard genes contained 20,000 baits and covered 158 target regions of 1,022,211 bp. The probe set was named Picea_hung_p1.0 (**Supplementary Data 1**, meta data in **Supplementary Table 2**).

The alignment of the probe set was tested on the reference genomes of five *Picea* species, *P. abies*^43^, *P. engelmannii*^2^, *P. engelmannii* × *P. glauca* × *P. sitchensis*^2^, *P. glauca*^2^, and *P. sitchensis*^2^, using minimap 2.22^44^.

A graphical abstract of the experimental and bioinformatic pipeline is presented in **Figure 2**.

**Figure 2.**
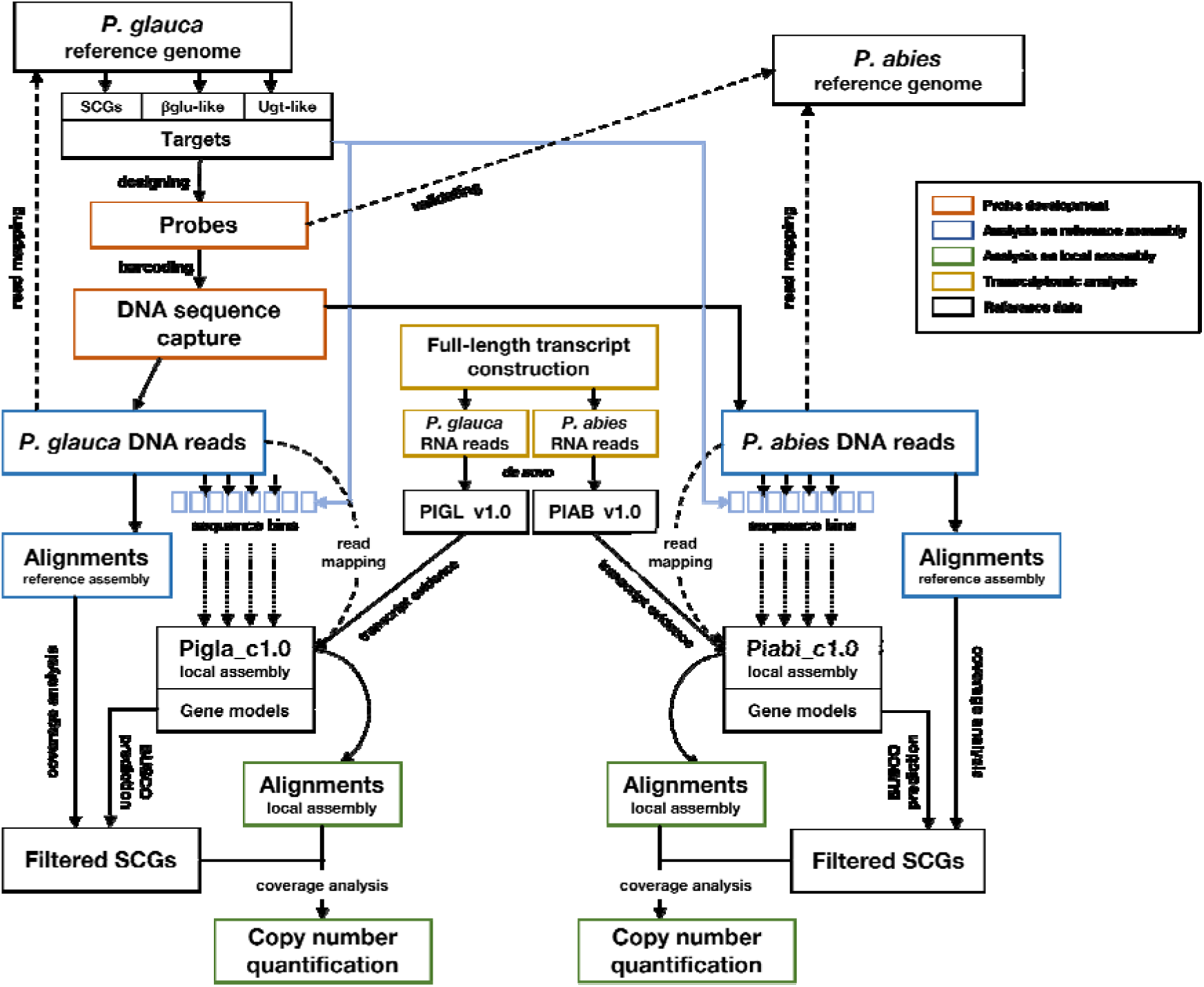
Experimental and bioinformatic pipeline of copy number quantification.

### Sequence capture and sequencing of genomic DNA

For each sample, 500 ng of genomic DNA was randomly fragmented to achieve a median size of ∼15 Kbp, using 1/8 reaction of NEBNext dsDNA Fragmentase (#M0348, New England Biolabs) incubated at 37°C for 5 minutes. Fragmented DNA was purified, end-repaired, dA-tailed, and ligated with adaptors for Illumina (#E6609, New England Biolabs) using NEBNext Ultra II Library Prep Kit for Illumina (#E7645, New England Biolabs). The sub-libraries were amplified and barcoded using a ProFlex PCR System (Thermo Fisher Scientific). The 25-µl reactions contained: 12.5 µl Kapa HiFi HotStart ReadyMix (Kapa Biosystems, United Kingdom), 2.5 µl NEBNext Multiplex Oligos for Illumina (10 µM) (#E6609, New England Biolabs), 10 µl adaptor-ligated DNA. The thermal cycling profile was: 94°C 3 min, 10 × [94°C 30 s, 65°C 4 min 30 s], 65°C 5 min, with ramp rate of each step at 3°C/s, to amplify fragments roughly from 1 to 10 Kbp. For all purification steps, a 0.4× AMPure XP (Beckman Coulter, United States) clean-up was used for size selection above ∼2 Kbp.

Hybridisation capture was performed with myBaits Custom 1–20K Kit (Daciel Arbor Biosciences, United States), using the Long Insert Protocol (manual version 5.02). The target-enriched sub-library was amplified using the same profiles as the barcoding amplification above, except using 25 cycles. All sub-libraries were pooled and normalised.

Nanopore libraries were constructed using the Ligation Sequencing Kit Chemistry 14 (SQK-LSK114, Oxford Nanopore Technologies, United Kingdom) using ∼200 fmol pooled library. Nanopore libraries were then sequenced on a R10.4.1 flow cell (FLO-PRO114M) at 400 bps (‘default’ mode) on a PromethION system (Oxford Nanopore Technologies) at the DNA Technologies & Expression Analysis Core Laboratory, UC Davis Genome Center.

All nanopore reads in this study were basecalled and demultiplexed from raw electrical signals using Guppy v6.0.0, trimmed for Nanopore and/or Illumina adaptors and split for chimeras using Porechop 0.2.4.

### Full-length cDNA (fl-cDNA) sequencing and transcriptome assembly

All total RNA samples were pooled to 1 µg for each species. First-strand synthesis, tailing, template switching, and extension were all completed with the SMARTer PCR cDNA Synthesis Kit (Takara Bio Europe, France). Full-length cDNA (fl-cDNA) library was constructed using the primers provided and the thermal cycling profile was: 94°C 3 min, 18× [94°C 30 s, 65°C 4 min 30 s], 65°C 5 min. For all purification steps, a 1.2× AMPure XP (Beckman Coulter, United States) clean-up was used. Nanopore libraries were constructed using the Ligation Sequencing Kit Chemistry 14 (SQK-LSK114, Oxford Nanopore Technologies) using ∼200 fmol pooled library. Nanopore libraries were then sequenced on a R10.3 flow cells (FLO-MIN111) on a GridION system (Oxford Nanopore Technologies, United Kingdom).

Fl-cDNA sequences were filtered for quality (Q score > 10) and trimmed for Nanopore adaptors and SMARTer PCR primers using Porechop 0.2.4^45^ with manual configuration. Filtered reads were used to construct the transcriptome using RNA-Bloom2^46^. Completeness of the transcriptomes were assessed using BUSCO v5.1.2^42^ with the embryophta_odb10 database. The transcriptome assemblies were named PIAB_v1 and PIGL_v1 respectively. Abundance of each transcript was quantified using minimap 2.22^44^ for mapping the raw fl-cDNA sequences to PIAB_v1 and PIGL_v1.

### Local sequence assembly and gene characterisation

To circumvent the extreme sequencing and computational requirements of whole-genome global assembly in spruce giga-genomes, local assembly could be used to assemble reads from the sequence capture and improve the characterisation of gene models ^47^, which may be incomplete in the reference genome.

Non-uniform sequence coverage due to the nature of sequence capture, rich repeat content, and biological variation had to be taken account into the local sequence assembly. For both species, all filtered reads were first searched against the gene models using BLAST+ and binned accordingly. Sequences in each bin were assembled using Canu 2.2^48^ with the parameters ‘maxInputCoverage=10000 corOutCoverage=10000 corMhapSensitivity=high corMinCoverage=0 redMemory=32 oeaMemory=32 batMemory=200’ according to the developers’ recommendations to retain as many reads as possible. The local sequence assemblies were named Piabi_c1.0 and Pigla_c1.0 respectively.

Gene models were characterised using both *ab initio* prediction and transcript evidence of PIAB_v1 and PIGL_v1 from the transcriptome assemblies above using MAKER 3.01.03^49^. The gene models were named Piabi_c1g and Pigla_c1g respectively.

### Gene analysis of ***β***glu-1 and Ugt5

Full-length cDNA sequences of *Pg*β*glu-1* (NCBI GenBank Accession: KJ780719.1)^12^, *PgUgt5* (KY963363.1), and *PgUgt5b* (KY963364.1)^14^ were searched against the Piabi_c1g and Pigla_c1g using BLAST 2.11.0+^41^. Homologous sequences of the cDNA and corresponding protein were subjected to local alignment with generalised affine gap costs (Altschul method) for β*glu-1* and *Ugt5* by using E-INS-i in MAFFT v7.490^50^ with 1,000 iterations. ModelTest-NG 0.17^51^ was used to select the best evolutionary model. A maximum likelihood phylogenetic tree of the cDNA sequences was built using RAxML-NG v. 1.0.2^52^ with the GTR+G4 substitution model with bootstrapping. The phylogenetic tree was midpoint-rooted and visualised using ggtree^53^, along with the protein alignment using ggmsa.

### Single-copy gene validation

The 139 putative single-copy genes were validated before use as internal standards. First, filtered genomic DNA reads were mapped to the reference genomes of *P. abies* and *P. glauca* (WS77111_v2) using minimap 2.22 with the specific option ‘-I 100g’ in both the indexing and mapping step given the large genome size. The alignment BAM files were converted to BED files using bam2bed 2.4.39 in the BEDOPS package^54^. Coverage statistics were calculated using the package TEQC^55^ in R. Normalised coverage for each target gene was calculated as the average coverage over the target bases divided by the average coverage over bases of all target genes. Second, BUSCO analysis was performed on the local gene models Piabi_c1.0g and Pigla_c1.0g to differentiate single-copy, duplicated, fragmented, and missing models.

### Copy number quantification

Most of the available CNV algorithms could only report relative copy number in forms of ratios^56^. We used single-copy genes as internal standards and were able to quantify absolute copy number by comparing coverage statistics of each gene model with that of the single-copy genes.

Filtered genomic DNA reads were mapped to the local assemblies Piabi_c1.0 and Pigla_c1.0. Normalisation of average coverage of the gene models was performed using the trimmed mean of M value (TMM), which accounted for factors of variations, such as sequencing depth, library composition, and gene length, with the edgeR package^57^. Copy numbers of β*glu-1* and *Ugt5b* were then calculated as their normalised counts divided by the geometric mean of the normalised counts of high-confidence single-copy genes. We only considered *Ugt5b* but not *Ugt5* because only *Ugt5b* was responsible for the synthesis of the biologically prevalent pungenin^14^. High-confidence single-copy genes were defined as those both within one standard deviation from mean for the log-normalised coverage in the reference assembly and classified as single-copy in the local assembly. CNV was detected by two-way ANOVA to test the significance of the effects of provenance, gene models, and their interaction on the total copy number.

### Validation of copy number quantification

We ran an independent copy number prediction with CNVPanelizer^58^ as parallel analysis to our copy number quantification using single-copy-gene reference. It was performed in *P. glauca* using a subsampling strategy similar to random forest for 10,000 replicates, and in particular compared the mean ratios of copy number of the complete gene form Pigla_c1g_00044 among provenances.

Quantitative PCR (qPCR) was also used to validate the copy number results and detection of CNV. The target amplicon started at exon 12 and ended at exon 13, which was unique in complete gene forms of β*glu-1*. The forward primer was 5’– CACAACCCCGCTTGAAGAAG–3’ and the reverse primer was 5’– TAACCTCGGACGTCTGCTCC–3’. The 10-µl reactions contained: 5 µl Luna Universal qPCR Master Mix (New England Biolabs), 0.25 µl forward primer (10 µM), 0.25 µl reverse primer (10 µM), and 4.5 µl of 20 ng genomic DNA. The thermal cycling profile was: 95°C 1 min, 45 × [95°C 15 s, 60°C 30 s with plate read]. The threshold cycle, known as C-half or C_1/2_, for each reaction was determined based on inflection point of the sigmoidal curve of the amplification profile. The amplicons were then validated by Sanger sequencing. Copy number quantification was calculated from a standard curve based on a serial dilution of a standard synthetic 2,000-bp oligomer of *Pg*β*glu-1* (Twist Bioscience, United States) which was conserved among all predicted gene forms.

## Results

### Performance of probe set and sequence capture

We developed a probe set targeting 158 genic sequences in *Picea glauca* including the sequences of the acetophenone biosynthesis genes *Pg*β*glu-1*, *PgUgt5*, and *PgUgt5b*, and 139 putative single-copy genes. The probes aligned well to the homologous sequences in the reference genomes of all five *Picea* species (**Supplementary Figure 1**). *P. glauca* had a 99.9% alignment rate, followed by *P. engelmannii* (98.13%), *P. abies* (97.79%), *P. sitchensis* (97.74%), and *P. engelmannii* × *P. glauca* × *P. sitchensis* (97.66%).

This newly designed probe set served for sequence capture of genomic DNA in individuals of *P. abies* (N = 38) and *P. glauca* (N = 40), which yielded a total 51.0 M reads with an N50 of 1,777 bp. The mean read number ± standard error (SE) for each sample was 357.49 K ± 16,397.46. The mean total sequence length ± SE was 551 ± 24.84 Mbp. The mean N50 ± SE was 1,691.79 ± 2.47 bp. We aligned the reads to the *P. abies* and *P. glauca* genomes and the mean alignment rate was 99.97%. The mean enrichment factor (target-to-background ratio) ± SE was 156.69 ± 4.64. The mean sensitivity ± SE was 69.88% ± 1.38%.

### Local sequence assembly and gene models

We produced local assemblies (Piabi_c1.0 and Pigla_c1.0) with the genomic sequence capture reads using a metagenomic approach. The two assemblies shared similar statistics: they contained 3,549 and 3,369 contigs and had a total read length of 9.28 and 9.11 Mbp, with a N50 of 2,644 and 2,726 bp, respectively. *In silico* gene prediction and transcript evidence of PIAB_v1 and PIGL_v1 produced 1,443 and 1,552 gene models (Piabi_c1.0g and Pigla_c1.0g) comprising of a total of 1.38 and 1.48 Mbp, with an average size of 957.5 and 951.7 bp, respectively. The statistics for the local genome assemblies, transcriptome assemblies, and the gene models can be found in **Supplementary Table 3**.

### Gene characterisation of ***β***glu-1 and Ugt5

Sequence similarity analysis of the gene models resulting from the local assemblies with the β*glu-1* model (KJ780719.1) gave 62 and 77 targets for *P. abies* and *P. glauca* respectively, which had substantial variation in terms of transcript abundance in the sample pool (**Figure 3**). A few of the gene models resembled the gene structure predicted in the β*glu-1* model (**Supplementary Figure 2**)^12^, which was located on the contig Pg-03r170320s1672760 and linkage group LG02 in the WS77111_v2 assembly of *P. glauca*^2^. For *P. abies*, Piabi_c1g_00049 and Piabi_c1g_00080 retained all 13 exons, while Piabi_c1g_00079 only missed the exon 10. For *P. glauca*, Pigla_c1g_00044 retained all 13 exons and Pigla_c1g_00123 had a split exon 7, while Pigla_c1g_00089 missed exon 10, Pigla_c1g_00279 had a truncated exon 13, and Pigla_c1g_00464 started with a truncated exon 4. All these complete or nearly complete gene models had considerably higher transcript levels compared to the other gene models, and transcripts in *P. glauca* were generally around 10-fold more abundant than those in *P. abies*.

**Figure 3.**
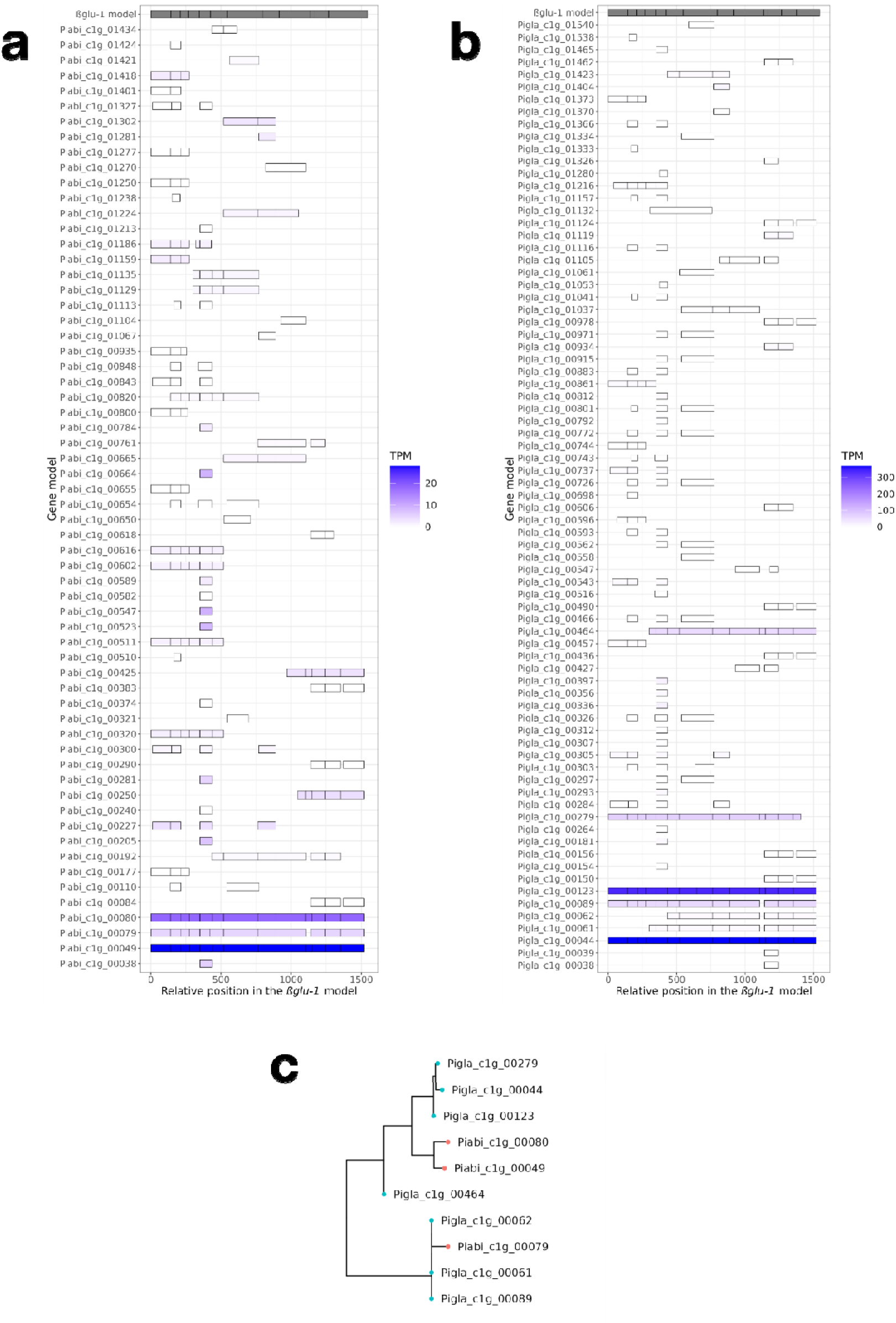
Exonic correspondence of βglu-1-like gene models in the local assemblies of **(a)** P. abies and **(b)** P. glauca against the reference βglu-1 gene model (KJ780719.1). The colour shows the transcript per million (TPM) of the fl-cDNA reads mapped against the gene models. **(c)** Reduced phylogenetic tree of the 10 complete and near-complete βglu-1 gene models in the local assemblies of P. abies (red tips) and P. glauca (blue tips). The complete phylogenetic tree of all 139 βglu-1-like gene models can be found in **Figure 6**.

Sequence similarity analysis of the gene models with the *Ugt5* (KY963363.1) and *Ugt5b* (KY963364.1) resulted in 2 and 33 targets for *P. abies* and 2 and 40 targets for *P. glauca* respectively, which had substantial variation in terms of transcript abundance (**Figure 4**). Many of the assembled sequences showed full-length or near-full-length correspondence with the *Ugt5b* model, which was located on the contig Pg-03r170320s1673621 and linkage group LG11 in the WS77111_v2 assembly of *P. glauca*. However, some of the *Ugt5b* models had much higher transcript abundance than other models, such as Piabi_c1g_00253, Piabi_c1g_00368, Piabi_c1g_00738, and Piabi_c1g_01026 in *P. abies*, and Pigla_c1g_00566 and Pigla_c1g_00787 in *P. glauca*.

**Figure 4.**
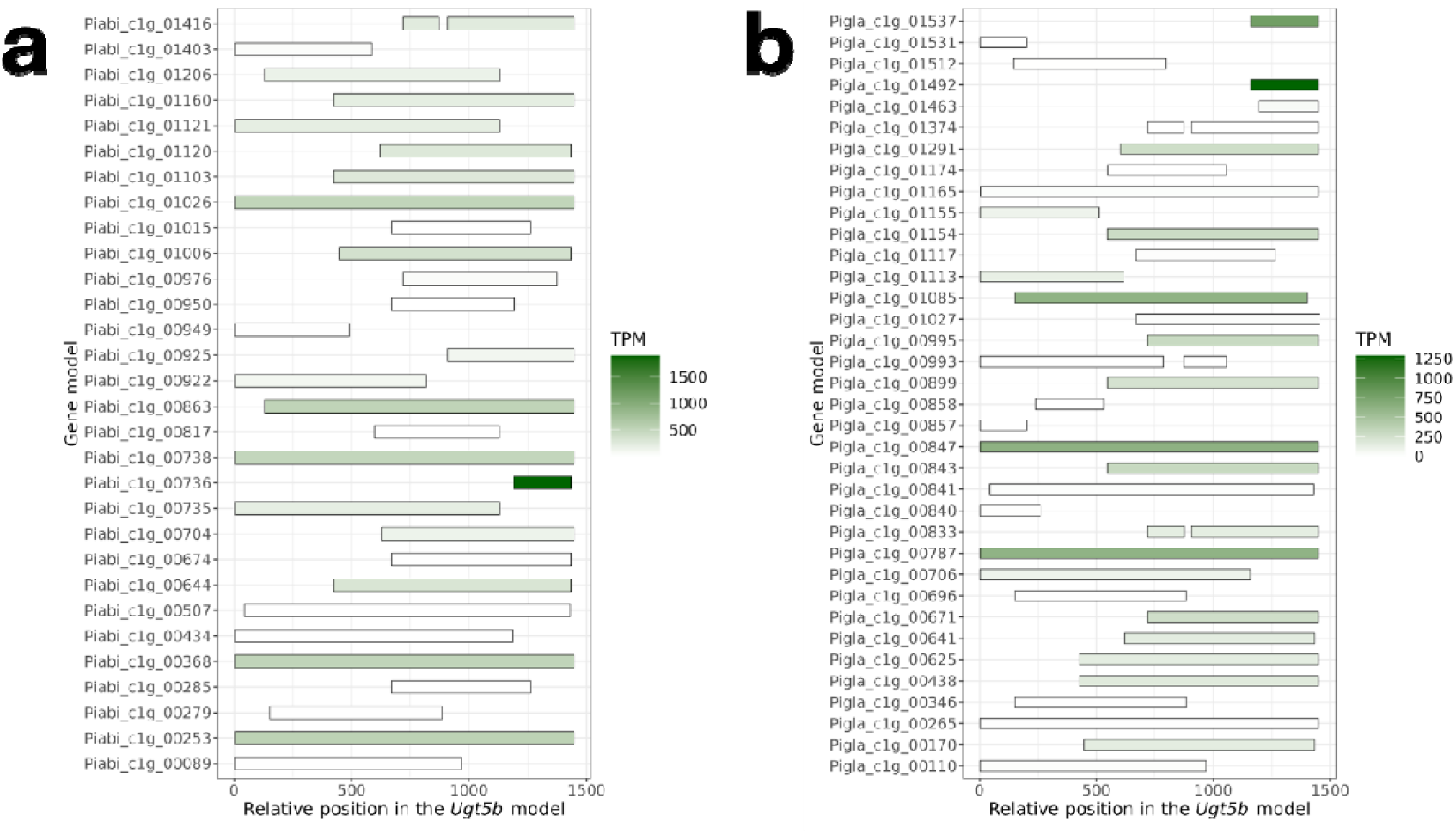
Exonic correspondence of Ugt5b-like gene models in the local assemblies of (a) P. abies and (b) P. glauca against the reference Ugt5b gene model (KY963364.1). The colour shows the transcript per million (TPM) of the fl-cDNA reads mapped against the gene models.

### Gene evolution of ***β***glu-1

We generated a rooted phylogenetic tree of 139 gene models that had a BLAST match (E < 1e–06) with the β*glu-1* model (KJ780719.1) to reconstruct the evolution of the β*glu-1*-like genes (**Supplementary Figure 3**). In particular, we investigated the complete β*glu-1* models which had 13 exons (**Figure 3c**), namely Piabi_c1g_00049, Piabi_c1g_00080 in *P. abies* and Pigla_c1g_00044 and Pigla_c1g_00123 in *P. glauca*. They formed a monophyletic clade in the reduced tree and a paraphyletic clade in the full tree, closely related to an ancestral Pigla_c1g_00464 that contained an exon 4 with a 5’-end truncation. All other gene models formed a separate cluster and were highly similar to each other as revealed by the near-zero branch lengths.

### Validation of single-copy genes

We analysed the putative single-copy gene set using cross-references from the reference and local assemblies with the BUSCO embryophyta_odb10 database as the standard. We observed that their log-normalised coverage in the reference genomes showed substantial variation in both species, with means of 1.31 and 1.06 and standard deviations of 0.84 and 0.65, respectively (**Figure 5**). On the other hand, we recovered 72 single-copy, 9 duplicated, and 14 fragmented genes in Piabi_c1g, and 79, 11, 11 respectively in Pigla_c1g. We retained 54 and 62 high-confidence single-copy genes in *P. abies* and *P. glauca*, respectively, which are within 1 standard deviation in the log-normalised coverage in the reference assembly and classified as complete in the local assembly. The two species shared 48 of these high-confidence single-copy genes (**Supplementary Table 4**).

**Figure 5.**
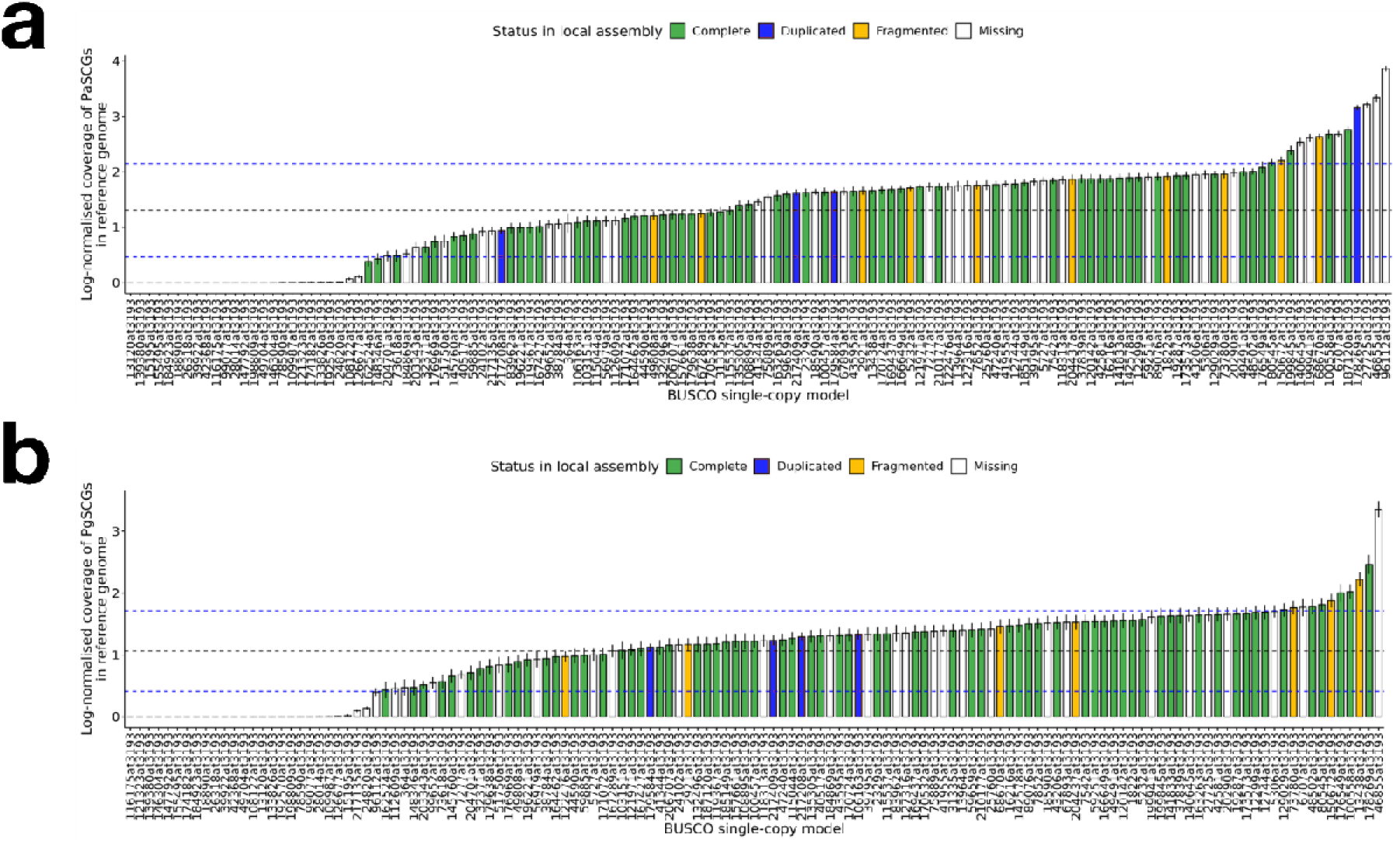
Log-normalised coverage of putative single-copy genes in the reference assemblies of **(a)** P. abies and **(b)** P. glauca. The genes are arranged in ascending order of their mean log-normalised coverage. The black and blue dashed lines indicate the mean ± 1 standard error. The colour of the bars represents their status in the local assemblies Piabi_c1.0 and Pigla_c1.0 respectively using BUSCO embryophyta_odb10 database as the reference.

### CNV of ***β***glu-1

We observed very high copy numbers and large CNV of β*glu-1* among populations of both *P. abies* and *P. glauca* (**Figure 6a and b**). The total copy number of all β*glu-1* gene models ranged between 56 copies in a *P. glauca* individual from the provenance 1338 (Twist Lake, Canada) and 381 copies in a *P. glauca* individual from the provenance 1987 (Woodlands County, Canada), representing a 6.8-fold difference between the two most extreme cases. In *P. abies*, the total copy number of β*glu-1* was significantly different among the gene models (Two-way ANOVA, *P* < 2e–16), but not among provenances (*P* = 0.552) and for their interaction (P = 0.282) (**Supplementary Table 5**). In *P. glauca*, it was significantly different among the provenances (*P* < 2e–16), among the gene models (*P* < 2e– 16), and also for their interaction (P = 0.000281) (**Supplementary Table 6**). Overall, the mean total copy number of β*glu-1* was higher in *P. glauca* than *P. abies*, which were 160 and 127 respectively (Wilcoxon rank sum exact test, *P* = 0.001535). The mean ratios of copy number of Pigla_c1g_00044 predicted using CNVPanelizer showed high congruence with our method using single-copy-gene references (*r* = 0.86, *P* = 2e–11, **Figure 6c and d**).

**Figure 6.**
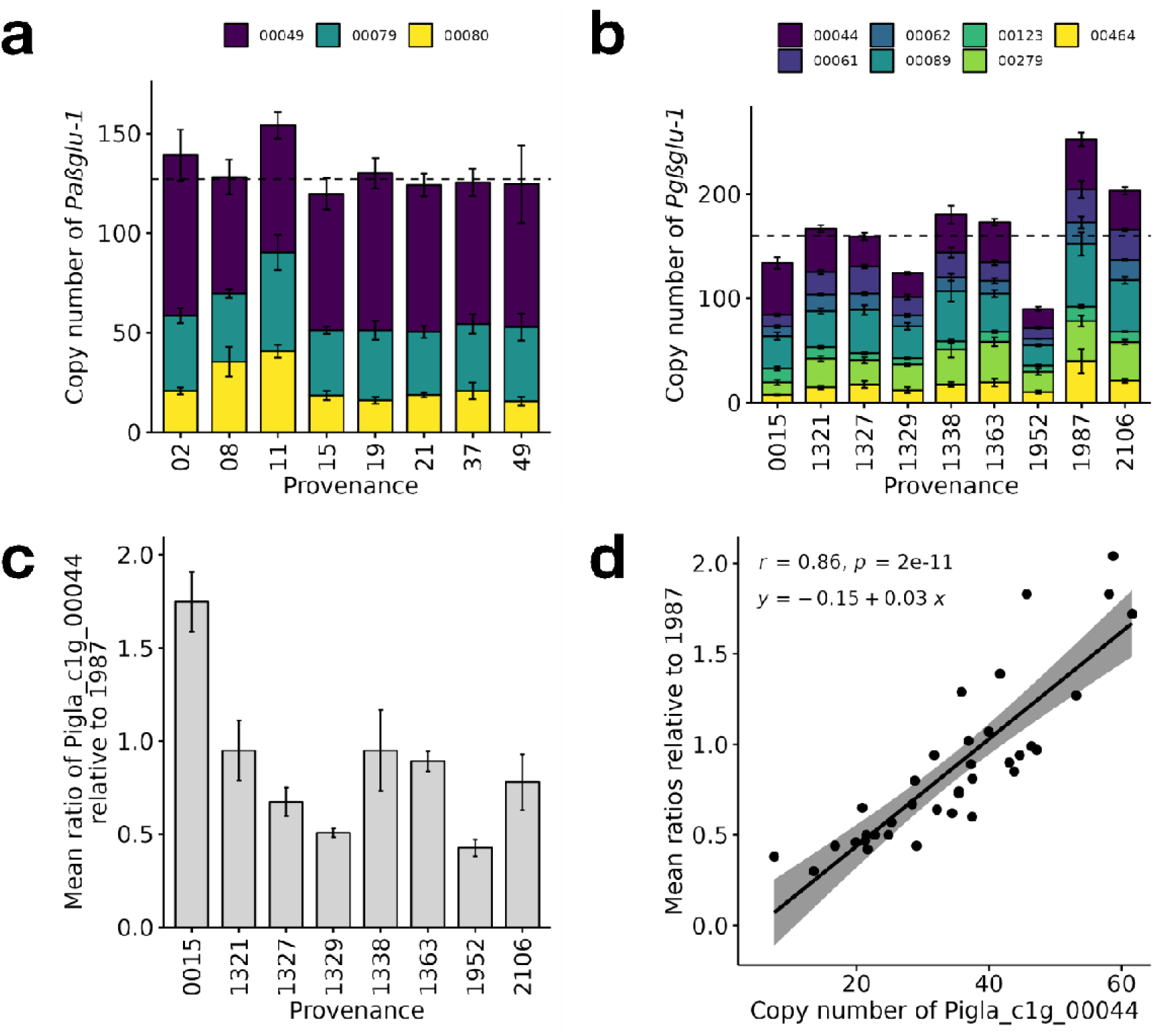
Copy number of complete and near-complete βglu-1 gene models determined by targeted sequencing in **(a)** P. abies and **(b)** P. glauca across their provenances. The error bar shows mean ± 1 standard error for each provenance. The dotted line shows the median of the total copy number. **(c)** Mean ratio of copy number of the complete gene form Pigla_c1g_00044 relative to provenance 1987 predicted using CNVPanelizer. **(d)** Correlation between the copy number of Pigla_c1g_00044 predicted using the single-copy-gene reference against the mean ratios predicted using CNVPanelizer.

We observed a similar scale of copy number and CNV of β*glu-1* in *P. glauca* using qPCR validation (**Supplementary Figure 4**). The copy number of β*glu-1* was significantly different among provenances (One-way ANOVA, *P* = 6.82e–5). The copy numbers were not significantly different between that assessed using sequence analysis and that using qPCR (*P* = 0.065, Welch two sample t-test). We did see discrepancies in the trends as generally fewer significant pairwise differences were found using qPCR than sequence analysis (**Supplementary Table 7** and **Supplementary Table 8**), for example, provenance 0015 was estimated to have a higher copy number using qPCR, but the general trend and difference for other provenances were similar to than found using sequence analysis.

### CNV of Ugt5b

In contrast to β*glu-1*, the *Ugt5b* gene models were detected as single-copy in most provenances in both species, with the medians of total copy number close to 1 (**Figure 7**) and no significant differences (Wilcoxon rank sum test, *P* = 0.135). However, there were a few clear outliers in both species. Specifically, the two *P. abies* provenances (08 and 11) had copy numbers of 14 and 12, respectively, and the two *P. glauca* provenances (1329 and 1952) had copy numbers of 11 and 4, respectively. The effects of provenance, gene model, and their interaction were all significant for both *P. abies* (*P* < 0.05) (**Supplementary Table 9**) and *P. glauca* (*P* < 0.05) (**Supplementary Table 10**).

**Figure 7.**
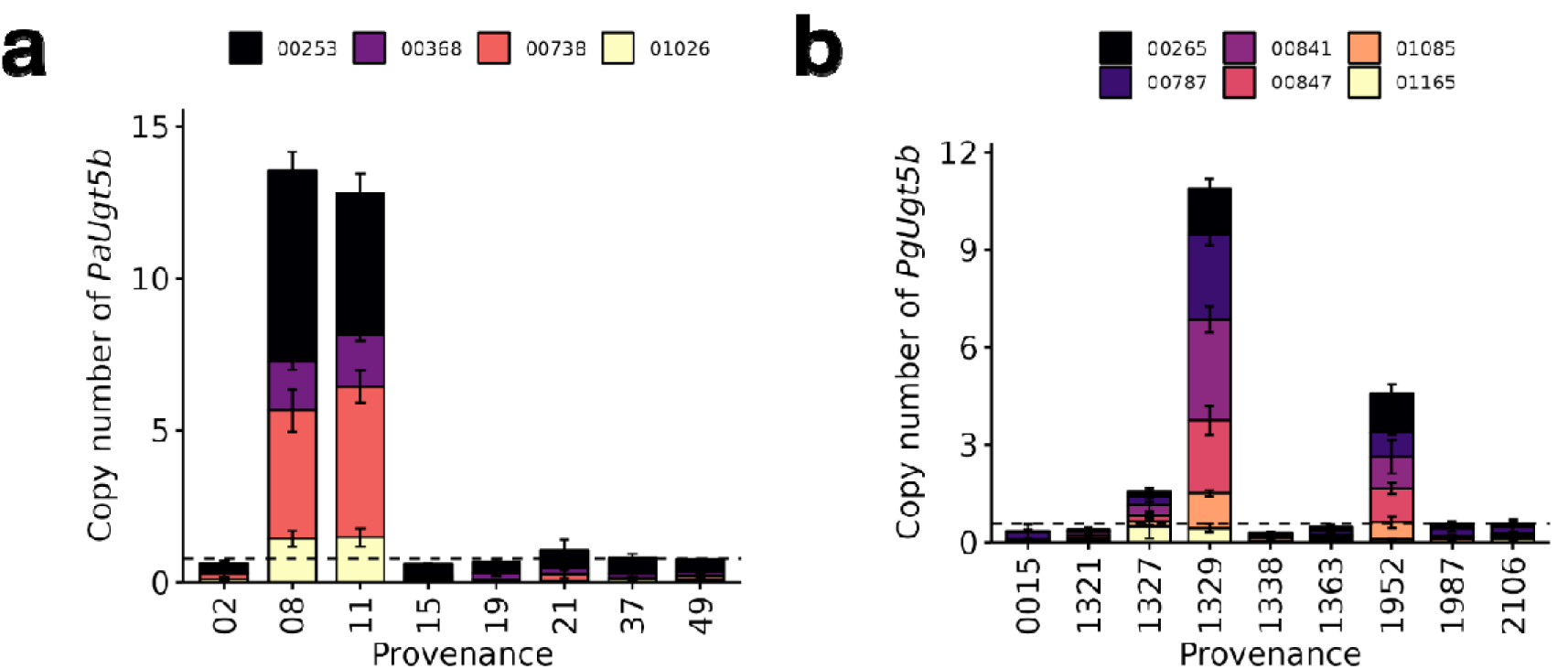
Copy number of complete and near-complete Ugt5b gene models in **(a)** P. abies and **(b)** P. glauca across their provenances.

## Discussion

### A novel method and probes for studying CNV in spruce gigagenomes

We have developed a long-insert target capture protocol to assess copy number of genes, which we optimised for a high throughput and quantitative comparison across samples. Most available copy number quantification algorithms have been developed for human disease diagnostics, particularly in cancers, where pairing of a control reference and an affected sample is the standard method of analysis^56,59^. A few recent studies have used coverage-based approaches for reference-free comparisons across samples and even populations^60,61^. Our method extended these approaches by incorporating probes that target single-copy genes, which serve as internal standards for copy number quantification of target genes in contrast to relative variations. We also implemented a bioinformatic pipeline incorporating local genome assembly for each target gene, which overcomes the biases from non-uniform coverage, as assembly algorithms tend to either collapse highly similar paralogues or exclude outlier-coverage reads^62^.

We assessed copy numbers of β*glu-1*-like and *Ugt5*-like genes in two keystone spruce species, *P. abies* and *P. glauca*. We also found near-perfect mapping rates of the probe set with other *Picea* spp. genomes. Therefore, the protocol is likely to be transferable to similar studies in any other conifer species, enabling large-scale, population-wide analyses that would be difficult to with other methods.

### Evolution and diversity of ***β***glu-1

We detected a total of five complete gene forms of β*glu-1* across the two species (Piabi_c1g_00049, Piabi_c1g_00080, Pigla_c1g_00044, Pigla_c1g_00123, Pigla_c1g_00279), which have all 13 exons and formed a paraphyletic clade. Therefore, our phylogenetic analysis suggests that an ancestral complete β*glu-1* gene emerged prior to the split of the *P. abies* and *P. glauca* lineages and only once, with gene duplication appearing to have occurred subsequently and independently within each lineage. One explanation for this observation is that the whole genome duplication in family Pinaceae around 200 to 342 million years ago, around the end-Permian extinction, may have generated new genes by sub- or neo-functionalisation^63^.

The absence of exon 1–5 (1–434) and 10 (369–380) in *Pg*β*glu-1* still yields gene transcripts. The structural model of the PgβGLU-1 enzyme predicted that the catalytic site residues E199 and E413 form hydrogen bonds with oxygen atoms O1 and O5 of the picein substrate, while residues T202, Y206, and R293 form hydrogen bonds with the phenolic moiety of picein^12^. While the near-complete gene forms retained these functionally important residues, it is likely that the missing exons impact the enzymatic activity of the encoded protein, and these shortened gene forms also have much lower expression than the complete forms. The absence of exon 1 (1–49) would also exclude a predicted N-terminal signal peptide (1–25), which is thought to target the PgβGLU-1 protein to the vacuole with release to the lumen after cleaving the hydrophobic signal peptide^12^. This predicted protein targeting is in general agreement with the histological observation of phenolic compounds in the vacuoles of mesophyll cells in spruce trees^64^.

The diversity of β*glu-1* gene copies discovered here was not captured in the most recent reference genome assemblies and gene model predictions of *P. abies*^43^ and *P. glauca*^2^, which only had one and two β*glu-1* gene models respectively. In *P. glauca*, the β*glu-1* gene models DB47_00003133 and DB47_00003134 had 10 and 6 exons respectively, implying errors in gene prediction and annotation. The genome of WS77111 v2 (*P. glauca*) has a BUSCO score of ∼50%, and its gene models only ∼20%, while that of the full-length transcriptome PIGL_v1 is ∼85%.

### Predicting single-copy genes in conifers

Surprisingly, we identified some very large copy numbers of up to 100 copies among genes that were assumed to be single-copy genes based on BUSCO prediction. This discrepancy between BUSCO prediction and our results may be explained by genomic difference between gymnosperms and other land plants. BUSCO is regarded as a reputable benchmarking tool to assess the completeness of genome assemblies and gene catalogues, with the assumption that a set of universal genes is expected to be found in a genome and in single-copy^65^. While the expectation of single-copy BUSCOs evolving under single-copy control has an evolutionary justification when the associated database (OrthoDB) was established^66^, none of the 61 species used to curate the embryophyte database until OrthoDB v10 is a gymnosperm^67^. Therefore, applying BUSCO predictions in gymnosperms would assume a conserved evolutionary history of these single-copy genes among gymnosperms and angiosperms, which is not supported by our results. Some of the main differences include the rate of molecular evolution in gymnosperms being seven times slower than in angiosperms^68^, Pinaceae having higher gene turnover rate than angiosperms^69^, fewer polyploid species among extant gymnosperms^63^, and highly distinct evolution of gene duplication and tandem-arrayed genes in conifers^70–75^.

### Rare extreme copy number and its variation of ***β***glu-1

The scale of up to 381 gene copies in a *P. glauca* individual and the level of variation we have observed in β*glu-1* are very rare in plants. The largest genic copy number known to date was 250–300 in a protein-coding sequencing encoding a tRNA ligase gene *RLG1a* in monkeyflower (*Mimulus guttatus*) from a population-pooled sequencing^60^, followed by 5– 160 copies of 5-enolpyruvylshikimate-3-phosphate synthase (*EPSPS*) in carelessweed *Amaranthus palmeri*^76^. In these examples, expansion of *EPSPS* conferred herbicide resistance via a gene dosage effect, and a triple copy of *RLG1a* was also associated with elevated *RLG1a* mRNA dosage but inconclusive for the *RLG1a* hundreds-copy extreme. In *P. glauca*, the conserved *R2R3-MYB* gene duplications may contribute to ∼100-fold increase in *PtMYB14* expression, which serves as important transcription factors in conifers, in the transgenic plantlets with a ubiquitous gene promoter than with a cinnamyl alcohol dehydrogenase tissue-preferential promoter^70^. Therefore, the contribution of CNV towards gene dosage could also be a plausible hypothesis for β*glu-1* with respect to insect resistance.

The mechanism underpinning extreme CNV remains poorly understood in plants. In the *EPSPS* case, cytogenetic analysis revealed that *EPSPS* clusters form extra-chromosomal circular DNA (eccDNA). These eccDNA are not integrated into a linear chromosome, are autonomously replicating, and transmittable to the next generations by tethering to mitotic and meiotic chromosomes^77^. Similarly, a tandem array of hundreds of *RLG1a* copies was unlinked to its single-copy locus, which leads to a working hypothesis that the cluster may also originate via rapid amplification in eccDNA and re-insertion into the nuclear genome^60^.

While eccDNAs may also provide a plausible mechanism to explain the emergence of large and variable copy numbers of β*glu-1* in *P. abies* and *P. glauca*, it remains a challenge to confirm. Many conifer genome assemblies are still highly fragmented and poorly annotated, due to their large size, long introns, and repetitivenes^69^. For example, ∼30% and ∼44% of the initially annotated genes in *P. abies* and loblolly pine (*Pinus taeda*) were fragmented or split into multiple scaffolds^43,78^. Therefore, it is challenging to align the extreme CNV of β*glu-1* in this study to the reference genomes and gene models, which may be collapsed during assembly, and to pinpoint their location. While conifer genome assemblies are expected to improve in the short future due to recent genomic advances, fluorescence *in situ* hybridisation probes mapping of β*glu-1* may be another approach to explore the origin of the very large number and variation of β*glu-1* copies.

### Among-provenance CNV in P. glauca but not in P. abies

CNV among provenances was only observed in *P. glauca* but not in *P. abies*. At the same time, the level of gene expression is on average more than 10-fold higher in *P. glauca* than in *P. abies*. Although fitness and survival are not directly tested in this study, these two findings indicate natural selection acting on *Picea* species.

The β*glu-1* gene contributes to resistance against spruce budworm *C. fumiferana* in *P. glauca*^10,11,16^, but *P. abies* is not affected by *C. fumiferana* in its natural species range^79^. Therefore, absence of CNV and low expression of β*glu-1* in *P. abies* may imply the absence of dosage effect of β*glu-1*, as it does not confer evolutionary advantage to the species. Meanwhile, low dosage of β*glu-1* may limit the autotoxicity and allelopathy of pungenol from foliage leachate, which could potentially decrease seed germination, in *P. abies*^80^. In contrast, the divergent evolution of copy number of β*glu-1* in *P. glauca* may correspond to the varying abundance and genetic structure of *C. fumiferana* across its distribution range^10,81^.

### The contrast of a very high copy number of ***β***glu-1 and single-copy Ugt5b

We analysed and observed large differences in gene copy numbers between *Ugt5b* and β*glu-1*, which encode enzymes that catalyse consecutive reactions in the spruce acetophenone pathway^12,14–16^. UGT5B glycosylates pungenol to form pungenin, and β*glu-1* releases piceol and pungenol from picein and pungenin, respectively. Tandem repeats of genes within the same pathway tend to possess a high degree of conservation in co-expression patterns, as found in the CNV in the benzylisoquinoline alkaloid pathway in opium poppy (*Papaver somniferum*)^82^, proteasome and ribosome pathways in *Arabidopsis thaliana*^83^, and duplicated genes in *Caenorhabditis elegans*^84^. Such a conserved co-expression may contribute to an increased flux through a metabolic pathway.

Perhaps surprisingly, we found that in both *P. abies and P. glauca*, the upstream gene *Ugt5b* is predominantly single-copy, while the downstream gene β*glu-1* is present with very high copy number. This implies that the acquisition and maintenance of additional copies in β*glu-1* does not effect a similar increase in copy number of *Ugt5b*. The two genes *Pg*β*glu-1* and *PgUgt5b* are located on different linkage groups (LG11 and LG02 respectively in *P. glauca*) and do not form operon-like gene clusters that might facilitate co-inheritance and co-regulation^85^. This is in contrast with some known gene clusters in plants which encode secondary metabolites for defence, such as benzoxazinoids, cyanogenic glucosides, terpenoids, and alkaloids^86^.

We propose three different hypotheses to explain the difference of copy number between genes that encode consecutive steps in the spruce acetophenone pathway. First, the two enzymes may have very different levels of enzymatic activity, including different substrate affinity and turnover, and thus the biochemical pathway may be well balanced despite a very different gene dosage. Second, it is possible that the majority of the β*glu-1* copies are not expressed, or their transcripts are not translated, and thus the effective dosage may be similar to that of *Ugt5b*. Third, it is possible that the two consecutive steps of the acetophenone pathway may not be acting in synchrony. The glucoside pungenin may accumulate slowly and constitutively over a period of time during the early summer based on the expression of a single-copy *Ugt5b*, while a larger gene dosage of β*glu-1* may be required later in mid-summer to release the defensive acetophenone, when the larval stage of spruce budworm is the most damaging^10^.

Besides the observation on the drastic difference in copy numbers, provenances 1329 and 1952 in *P. glauca* have higher copy numbers of *PgUgt5b* than other provenances, yet lower copy numbers of *Pg*β*glu-1*. This may imply an accumulation of glucosidic forms without effective synthesis of the acetophenones, and thus may lead to higher susceptibility in these provenances. However, without gene expression and metabolic data, we caution that we are not able to draw a conclusive relationship between the genotypes and phenotypes.

### Implications for genomic research in conifers

Our study provides a detailed characterisation of the gene forms, their sequences, and CNVs of a beta-glucosidase gene β*glu-1* in *P. abies* and *P. glauca*, with implications for understanding insect resistance in keystone spruce species and the evolution of complex conifer genomes. We generate original hypotheses for testing the emergence and mechanism of large and variable copy numbers, the potential dosage effect on phenotype, and the varying copy number of other genes within the same pathway. The next steps along this line of research would thus involve testing relationship between the absolute copy number of these defence-related genes, their expression, and the products using a bigger sample size and wider geographic coverage.

The challenge of incomplete and uncertain genome assemblies, despite advances in sequencing technologies, is likely to continue for conifer gigagenomes. We have developed experimental and bioinformatic approaches that overcome the limits of many current genome assemblies, and also developed new resources for genomic research in conifers.

## Declarations

### Ethics approval and consent to participate

Not applicable.

### Consent for publication

Not applicable.

### Availability of data and materials

Sequence data generated in this project, including the raw reads of genomic sequence capture and full-length transcriptome sequencing, local genome assemblies (Piabi_c1.0 and Pigla_c1.0), and full-length transcriptome assemblies (PIAB_v1 and PIGL_v1), can be publicly accessed in NCBI GenBank under the BioProject PRJNA940966 (http://www.ncbi.nlm.nih.gov/bioproject/940966). BioSamples are also registered under this BioProject.

### Authors’ contributions

T.H.H.: designed the study, processed the DNA samples, conceived and conducted the experimental and bioinformatic analyses, drafted the manuscript, and secured funding for the project;

E.T.Y.W.: processed the RNA samples, and revised the manuscript;

P.Z.: collected the samples and revised the manuscript;

A.J.: collected the samples, revised the manuscript, and secured funding for the project;

A.U.: collected the samples and revised the manuscript;

N.E.: collected the samples, revised the manuscript, and secured funding for the project;

J.Boh.: developed and provided the initial genomic data for *P. glauca*, and revised the manuscript;

J.Bou.: developed and provided the initial genomic data for *P. glauca*, and revised the manuscript;

I.B.: developed and provided the initial genomic data for *P. glauca*, and revised the manuscript;

S.M.C.: supervised the study, revised the manuscript, and secured funding for the project;

J.J.M.: designed and supervised the study, revised the manuscript, and secured funding for the project.

## Acknowledgements

Not applicable.

