## Supplementary Information for "Long-insert sequence capture detects high copy numbers in a defence-related beta-glucosidase gene β*glu-1* with large variations in white spruce but not Norway spruce"

Tin Hang Hung^1,*^
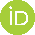
<https://orcid.org/0000-0001-9853-2053>, Ernest T. Y. Wu^1^, Pauls Zeltiņš^2^
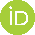
<https://orcid.org/0000-0002-6286-5814>, Āris Jansons^2^
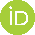
<https://orcid.org/0000-0001-7981-4346>, Aziz Ullah^3^
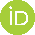
<https://orcid.org/0000-0001-6155-8874>, Nadir Erbilgin^3^
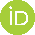
<https://orcid.org/0000-0001-9912-8095>, Joerg Bohlmann^4,5,6^
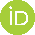
<https://orcid.org/0000-0002-3637-7956>, Jean Bousquet^7^, Inanc Birol^8^
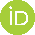
<https://orcid.org/0000-0003-0950-7839>, Sonya M. Clegg^1^
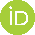
^<https://orcid.org/0000-0002-3092-3864>^, John J. MacKay^1,*^
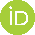
<https://orcid.org/0000-0002-4883-195X>

1. Department of Biology, University of Oxford, Oxford OX1 3RB, United Kingdom
2. Latvian State Forest Research Institute “Silava”, LV-2169 Salaspils, Latvia
3. Department of Renewable Resources, University of Alberta, Edmonton, AB, T6G 2E3, Canada
4. Michael Smith Laboratories, University of British Columbia, Vancouver, BC, V6T 1Z4, Canada
5. Department of Botany, University of British Columbia, Vancouver, BC, V6T 1Z4, Canada
6. Department of Forest and Conservation Sciences, University of British Columbia, Vancouver, BC, V6T 1Z4, Canada
7. Canada Research Chair in Forest Genomics, Forest Research Centre, Université Laval, Québec, QC, G1V 0A6, Canada
8. Canada’s Michael Smith Genome Sciences Centre, Vancouver, BC, V5Z 4S6, Canada

Corresponding authors:

*T.H.H.:; *J.J.M.:

**Supplementary Table 1.** Provenances of the samples analysed in this study for *P. abies* and *P. glauca*.

| Species | Provenance | Geographical location | | | Elevation | Latitude | Longitude | # Samples |
| --- | --- | --- | --- | --- | --- | --- | --- | --- |
| *P. abies* (Pa) | 02 | Slovakia | Tatra mountains | Smolnick | 700 | 48.76667 | 20.76667 | 5 |
|  | 08 | Denmark | Hasede |  | 20 | 55.75 | 12.2 | 3 |
|  | 11 | Romania | Carpathian mountains | Falcau Levinsen | 850 | 47.9 | 25.46667 | 5 |
|  | 15 | Germany | Harz and foothills | Babben | 100 | 51.7 | 13.78333 | 5 |
|  | 19 | Germany | Böhmerwald | Reinhardtsdorf | 210 | 50.88333 | 14.16667 | 5 |
|  | 21 | Germany | Northwest Europe | Grünhaus | 100 | 49.33333 | 8.583333 | 5 |
|  | 37 | Latvia | Kuldīga |  | 60 | 57 | 22.3 | 5 |
|  | 49 | Norway | Telemark |  | 250 | 59.5 | 8.8 | 5 |
| *P. glauca* (Pg) | 0015 | Canada | Alberta | Slave Lake Forest | 731 | 55.27221 | -114.78 | 4 |
|  | 1321 | Canada | New Brunswick | Upper Green River | 304 | 47.53863 | -68.2139 | 5 |
|  | 1327 | Canada | Quebec | Dasserat Twp | 289 | 48.16793 | -79.4152 | 5 |
|  | 1329 | Canada | Quebec | Cimon Twp | 198 | 45.8999 | -75.0985 | 4 |
|  | 1338 | Canada | Ontario | Twist Lake | 425 | 49.02692 | -93.075 | 5 |
|  | 1363 | Canada | Saskatchewan | Old Channel Riv | 266 | 53.93641 | -102.62 | 5 |
|  | 1952 | Canada | Alberta | Fox Creek | 881 | 54.39879 | -116.808 | 4 |
|  | 1987 | Canada | Alberta | Woodlands County | 870 | 54.19958 | -115.638 | 4 |
|  | 2106 | Canada | Alberta | Lesser Slave Riv | 580 | 55.30458 | -114.836 | 4 |

**Supplementary Table 2.** Details of the genomic regions targeted by the sequence capture bait set Picea_hung_p1.0.

*See separate spreadsheet*

**Supplementary Table 3.** Statistics of local genome assembly, transcriptome assembly, and gene models of *P. abies* and *P. glauca*.

|  | Piabi_c1.0 | Pigla_c1.0 | PIAB_v1 | PIGL_v1 | Piabi_c1.0g | Pigla_c1.0g |
| --- | --- | --- | --- | --- | --- | --- |
| Number of sequences | 3,549 | 3,369 | 137,487 | 128,738 | 1,443 | 1,552 |
| Total length of sequences | 9,275,211 | 9,116,455 | 160,667,657 | 278,747,468 | 1,381,689 | 1,477,113 |
| Minimum length | 1,079 | 1,049 | 200 | 200 | 21 | 15 |
| Average length | 2,613.5 | 2,706 | 1,168.6 | 2,165.2 | 957.5 | 951.7 |
| Maximum length | 18,952 | 18,909 | 11,633 | 23,709 | 11,580 | 14,935 |
| N50 | 2,644 | 2,726 | 1,527 | 3,020 | 1,400 | 1,390 |

**Supplementary Figure 1.** Alignment rates of the probe set Picea_hung_p1.0 on the reference genomes of 5 *Picea* species.

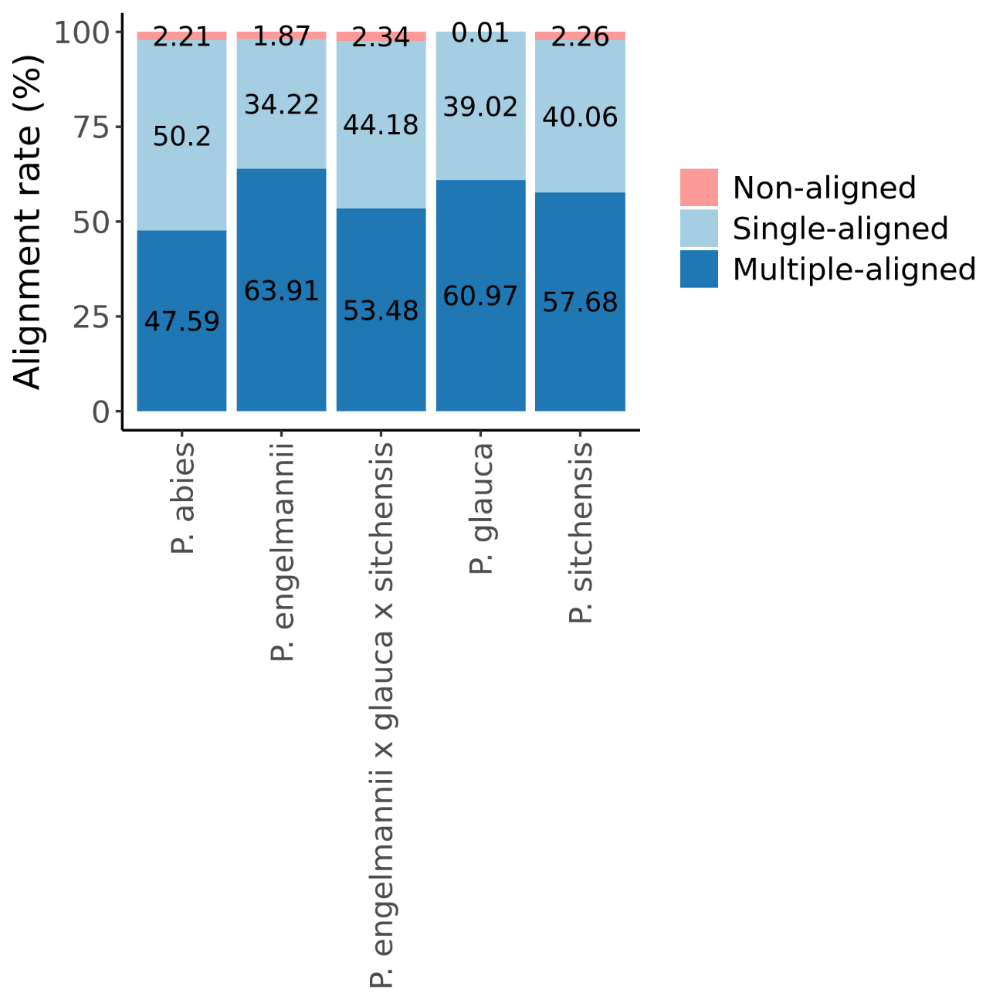

**Supplementary Figure 2.** Syntenic relationship of 7 complete or near-complete *Pgβglu-1* gene forms in *P. glauca* with the reference *Pgβglu-1* gene model (KJ780719.1).

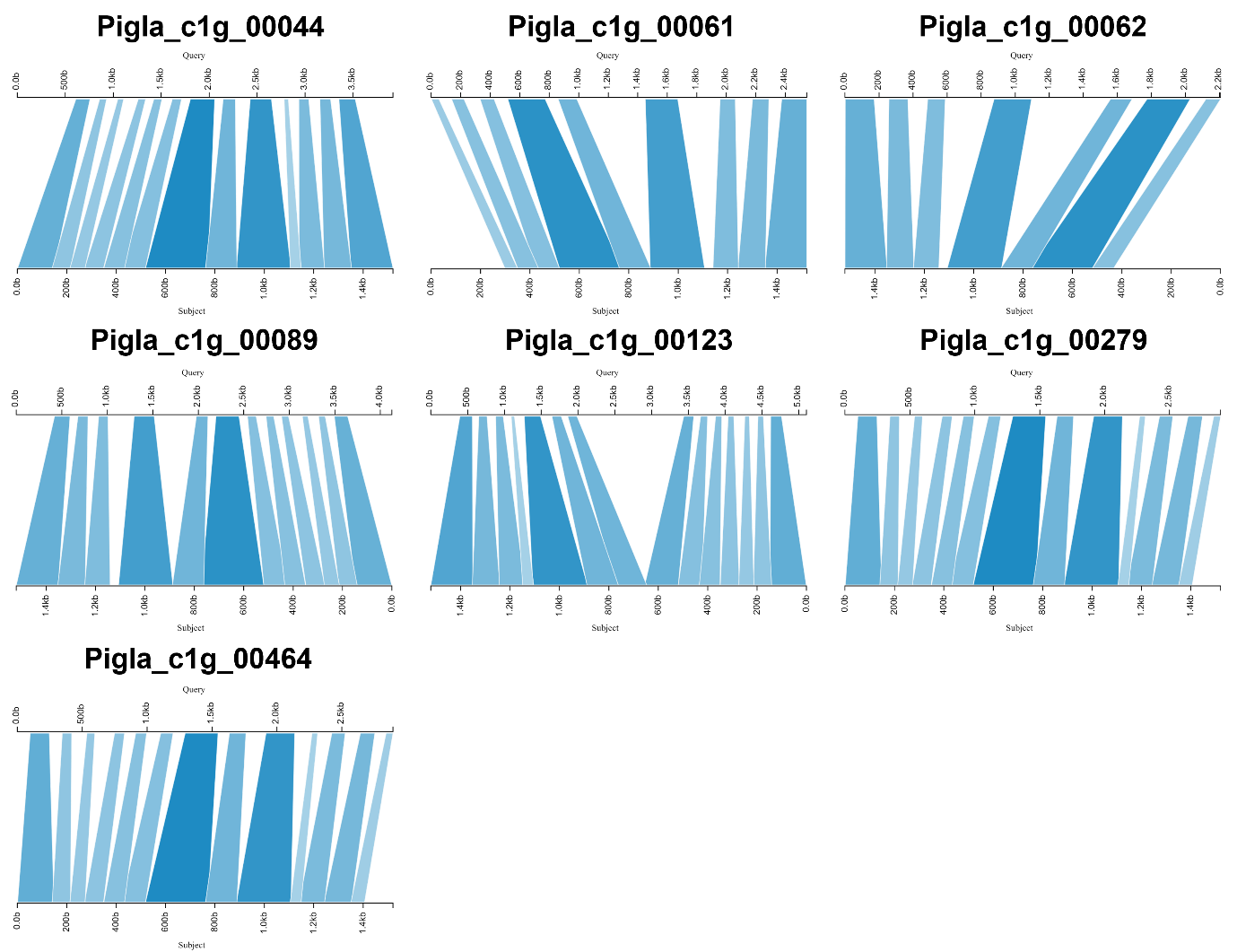

**Supplementary Figure 3.** Phylogenetic tree of 139 *βglu-1*-like gene models in the local assemblies of *P. abies* (red tips) and *P. glauca* (blue tips) and their protein multiple sequence alignment. The bold tip labels denote the complete and near-complete gene forms of *βglu-1*.

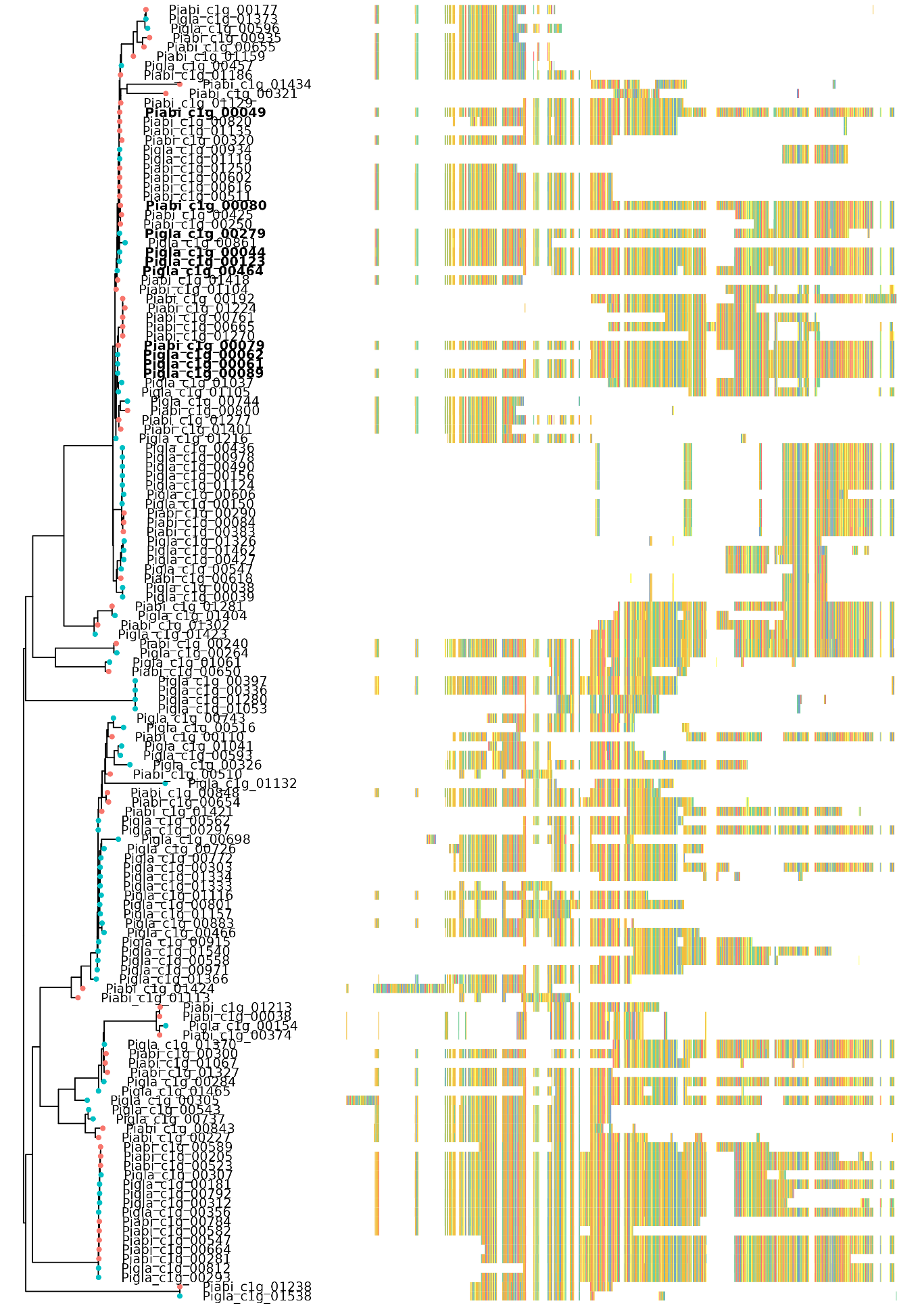

**Supplementary Table 4.** Details of the high-confidence single-copy genes shared by *P. abies* and *P. glauca*.

*See separate spreadsheet*

**Supplementary Table 5.** Analysis of variance (ANOVA) table of the total copy number of *Paβglu-1* in *P. abies*.

| **Factor** | **df** | **Sum sq** | **Mean sq** | **P-value** |  |
| --- | --- | --- | --- | --- | --- |
| Provenance | 7 | 1419 | 203 | 0.552 |  |
| Gene models | 2 | 47993 | 23997 | <2e–16 | *** |
| Interaction | 14 | 4057 | 290 | 0.282 |  |
| Residuals | 90 | 21544 | 239 |  |  |

**Supplementary Table 6.** Analysis of variance (ANOVA) table of the total copy number of *Pgβglu-1* in *P. glauca*.

| **Factor** | **df** | **Sum sq** | **Mean sq** | **P-value** |  |
| --- | --- | --- | --- | --- | --- |
| Provenance | 8 | 10163 | 1270 | <2e–16 | *** |
| Gene models | 6 | 30305 | 5051 | <2e–16 | *** |
| Interaction | 48 | 7060 | 147 | 0.000281 | *** |
| Residuals | 217 | 15594 | 72 |  |  |

**Supplementary Figure 4.** (a) Standard curve of *Pgβglu-1* synthetic oligomer. (b) Copy number of *Pgβglu-1* in *P. glauca* across their provenances estimated by qPCR. The error bar shows mean ± 1 standard error.

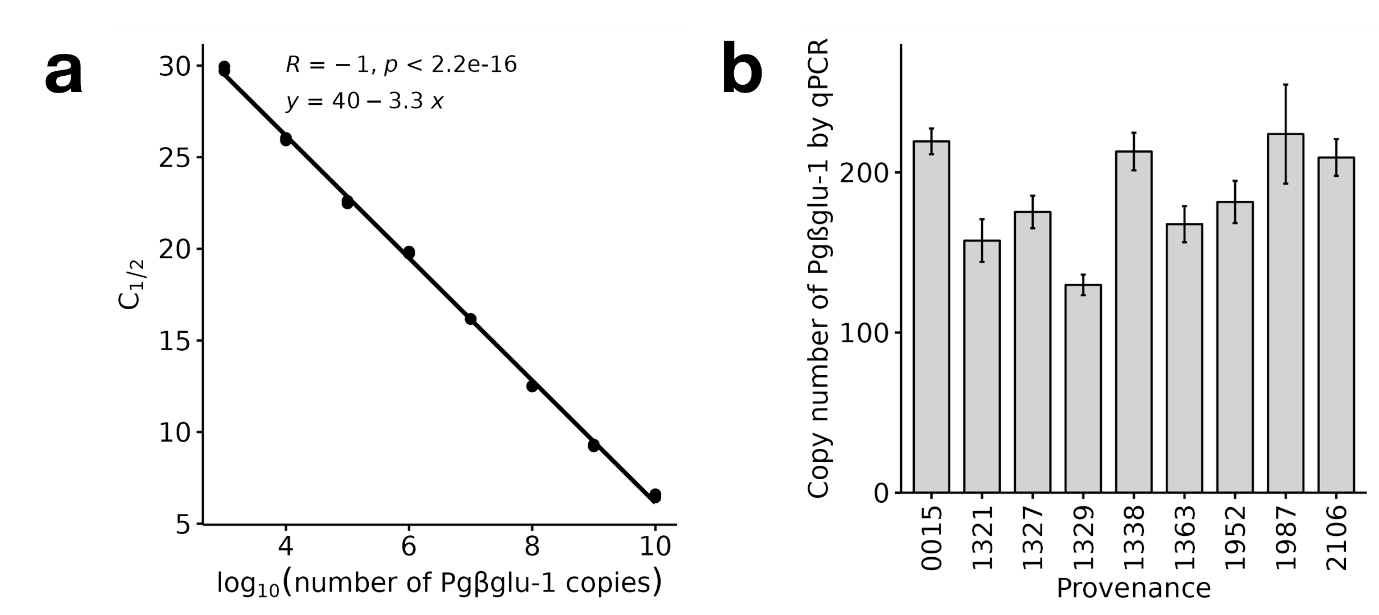

**Supplementary Table 7.** TukeyHSD pairwise comparison table of the total copy number of *Pgβglu-1* in *P. glauca* estimated with sequence analysis.

|  | **Difference** | **Lower** | **Upper** | **Adjusted** |  |
| --- | --- | --- | --- | --- | --- |
| 1321-0015 | 4.704126 | -2.03053 | 11.43878 | 0.416799 |  |
| 1327-0015 | 3.634558 | -3.10009 | 10.36921 | 0.751474 |  |
| 1329-0015 | -1.38341 | -8.48235 | 5.71554 | 0.999539 |  |
| 1338-0015 | 6.636293 | -0.09836 | 13.37095 | 0.056936 |  |
| 1363-0015 | 5.591683 | -1.14297 | 12.32634 | 0.192034 |  |
| 1952-0015 | -6.26477 | -13.3637 | 0.834173 | 0.13205 |  |
| 1987-0015 | 16.94709 | 9.848143 | 24.04604 | 0 | *** |
| 2106-0015 | 9.895333 | 2.796386 | 16.99428 | 0.000648 |  |
| 1327-1321 | -1.06957 | -7.41906 | 5.279924 | 0.999845 |  |
| 1329-1321 | -6.08753 | -12.8222 | 0.64712 | 0.112067 |  |
| 1338-1321 | 1.932167 | -4.41732 | 8.281659 | 0.98942 |  |
| 1363-1321 | 0.887557 | -5.46193 | 7.237048 | 0.999963 |  |
| 1952-1321 | -10.9689 | -17.7036 | -4.23425 | 2.55E-05 | *** |
| 1987-1321 | 12.24296 | 5.508312 | 18.97762 | 1.4E-06 | *** |
| 2106-1321 | 5.191207 | -1.54345 | 11.92586 | 0.281022 |  |
| 1329-1327 | -5.01797 | -11.7526 | 1.716687 | 0.326186 |  |
| 1338-1327 | 3.001735 | -3.34776 | 9.351226 | 0.863159 |  |
| 1363-1327 | 1.957124 | -4.39237 | 8.306616 | 0.988492 |  |
| 1952-1327 | -9.89933 | -16.634 | -3.16468 | 0.000238 | *** |
| 1987-1327 | 13.31253 | 6.577879 | 20.04718 | 1E-07 | *** |
| 2106-1327 | 6.260775 | -0.47388 | 12.99543 | 0.091307 |  |
| 1338-1329 | 8.0197 | 1.285047 | 14.75435 | 0.007369 | ** |
| 1363-1329 | 6.97509 | 0.240437 | 13.70974 | 0.036018 | * |
| 1952-1329 | -4.88137 | -11.9803 | 2.21758 | 0.439283 |  |
| 1987-1329 | 18.3305 | 11.23155 | 25.42944 | 0 | *** |
| 2106-1329 | 11.27874 | 4.179793 | 18.37769 | 4.55E-05 | *** |
| 1363-1338 | -1.04461 | -7.3941 | 5.304881 | 0.99987 |  |
| 1952-1338 | -12.9011 | -19.6357 | -6.16641 | 3E-07 | *** |
| 1987-1338 | 10.3108 | 3.576145 | 17.04545 | 0.000103 | *** |
| 2106-1338 | 3.25904 | -3.47561 | 9.993693 | 0.846904 |  |
| 1952-1363 | -11.8565 | -18.5911 | -5.1218 | 3.5E-06 | *** |
| 1987-1363 | 11.35541 | 4.620755 | 18.09006 | 1.08E-05 | *** |
| 2106-1363 | 4.30365 | -2.431 | 11.0383 | 0.543453 |  |
| 1987-1952 | 23.21187 | 16.11292 | 30.31081 | 0 | *** |
| 2106-1952 | 16.16011 | 9.06116 | 23.25905 | 0 | *** |
| 2106-1987 | -7.05176 | -14.1507 | 0.04719 | 0.053062 |  |

**Supplementary Table 8.** TukeyHSD pairwise comparison table of the total copy number of *Pgβglu-1* in *P. glauca* estimated with qPCR.

|  | **Difference** | **Lower** | **Upper** | **Adjusted** |  |
| --- | --- | --- | --- | --- | --- |
| 1321-0015 | -61.97411 | -125.871155 | 1.922936 | 0.0643039 |  |
| 1327-0015 | -44.08542 | -107.982465 | 19.811626 | 0.4123433 |  |
| 1329-0015 | -89.655214 | -157.008614 | -22.301814 | 0.0019283 | ** |
| 1338-0015 | -6.326815 | -70.223861 | 57.57023 | 0.9999966 |  |
| 1363-0015 | -51.707074 | -115.604119 | 12.189972 | 0.2097686 |  |
| 1952-0015 | -37.842613 | -105.196012 | 29.510787 | 0.6835247 |  |
| 1987-0015 | 4.551799 | -62.801601 | 71.905199 | 0.9999998 |  |
| 2106-0015 | -10.0717 | -77.4251 | 57.2817 | 0.9999189 |  |
| 1327-1321 | 17.88869 | -42.354022 | 78.131402 | 0.9890056 |  |
| 1329-1321 | -27.681104 | -91.57815 | 36.215941 | 0.8994142 |  |
| 1338-1321 | 55.647294 | -4.595418 | 115.890006 | 0.09297 |  |
| 1363-1321 | 10.267036 | -49.975676 | 70.509748 | 0.9997827 |  |
| 1952-1321 | 24.131497 | -39.765548 | 88.028542 | 0.9520851 |  |
| 1987-1321 | 66.525909 | 2.628863 | 130.422954 | 0.0349648 | * |
| 2106-1321 | 51.90241 | -11.994636 | 115.799455 | 0.2056696 |  |
| 1329-1327 | -45.569794 | -109.46684 | 18.327251 | 0.3669219 |  |
| 1338-1327 | 37.758604 | -22.484108 | 98.001317 | 0.5449608 |  |
| 1363-1327 | -7.621654 | -67.864366 | 52.621058 | 0.9999774 |  |
| 1952-1327 | 6.242807 | -57.654238 | 70.139853 | 0.999997 |  |
| 1987-1327 | 48.637219 | -15.259827 | 112.534264 | 0.2816677 |  |
| 2106-1327 | 34.01372 | -29.883326 | 97.910765 | 0.742602 |  |
| 1338-1329 | 83.328398 | 19.431353 | 147.225444 | 0.0025868 | ** |
| 1363-1329 | 37.94814 | -25.948905 | 101.845186 | 0.6159102 |  |
| 1952-1329 | 51.812601 | -15.540799 | 119.166001 | 0.2685939 |  |
| 1987-1329 | 94.207013 | 26.853613 | 161.560413 | 0.0009085 | *** |
| 2106-1329 | 79.583514 | 12.230114 | 146.936914 | 0.0092618 | ** |
| 1363-1338 | -45.380258 | -105.62297 | 14.862454 | 0.2948001 |  |
| 1952-1338 | -31.515797 | -95.412843 | 32.381248 | 0.8132893 |  |
| 1987-1338 | 10.878614 | -53.018431 | 74.77566 | 0.9997843 |  |
| 2106-1338 | -3.744885 | -67.64193 | 60.152161 | 0.9999999 |  |
| 1952-1363 | 13.864461 | -50.032585 | 77.761506 | 0.9987231 |  |
| 1987-1363 | 56.258872 | -7.638173 | 120.155918 | 0.1286628 |  |
| 2106-1363 | 41.635374 | -22.261672 | 105.532419 | 0.4917278 |  |
| 1987-1952 | 42.394412 | -24.958988 | 109.747811 | 0.5392263 |  |
| 2106-1952 | 27.770913 | -39.582487 | 95.124312 | 0.9222559 |  |
| 2106-1987 | -14.623499 | -81.976899 | 52.729901 | 0.9987174 |  |

**Supplementary Table 9.** Analysis of variance (ANOVA) table of the total copy number of *PaUgt5b* in *P. abies*.

| **Factor** | **df** | **Sum sq** | **Mean sq** | **P-value** |  |
| --- | --- | --- | --- | --- | --- |
| Provenance | 7 | 240.38 | 34.34 | <2e–16 | *** |
| Gene models | 3 | 30.74 | 10.25 | <2e–16 | *** |
| Interaction | 21 | 72.12 | 3.43 | <2e–16 | *** |
| Residuals | 120 | 27.78 | 0.23 |  |  |

**Supplementary Table 10.** Analysis of variance (ANOVA) table of the total copy number of *PgUgt5* in *P. glauca*.

| **Factor** | **df** | **Sum sq** | **Mean sq** | **P-value** |  |
| --- | --- | --- | --- | --- | --- |
| Provenance | 8 | 68.20 | 8.525 | <2e–16 | *** |
| Gene models | 5 | 3.61 | 0.722 | 5.91e–6 | *** |
| Interaction | 40 | 21.03 | 0.526 | 1.52e–14 | *** |
| Residuals | 186 | 19.39 | 0.104 |  |  |
